# PCR-free, label-free detection of sequence-specific DNA with single-molecule sensitivity using *in vitro* N-hybrid system in microfluidic drops

**DOI:** 10.1101/2022.01.09.475569

**Authors:** Yizhe Zhang, David A. Weitz

## Abstract

We propose a novel method that can detect DNA with high specificity at the single-molecule level by employing the *in vitro* N-hybrid strategy realized in sub-picoliter microfluidic drops. It detects target DNA based on the specific interactions of the target-encoded proteins with their partner molecules, and achieves single-molecule sensitivity via signal-transduction and signal-amplification during gene-expression processes in a sub-picoliter droplet, therefore effectively avoiding complicated procedures in labeling-based methods or biases and artifacts in PCR-based methods.

## Introduction

Biological studies on single-molecule level can help us to elucidate the details in cellular behaviors that might otherwise be buried in ensemble measurements (1). A large number of breakthroughs in fundamental research such as DNA folding kinetics (2–4), protein-DNA interactions (5–7), DNA replication (8–14) and molecular transport (15, 16), have benefited from the development in single molecule techniques. However, most of the prevalent approaches in single-molecule detection to date, for example, fluorescence/Förster resonance energy transfer (FRET), fluorescence correlation spectroscopy (FCS), total internal reflection fluorescence microscopy (TIRF) and surface-enhanced Raman spectroscopy (SERS), require the labeling of the target (17, 18), which might interfere with the properties of interest and make the scarce targets unrecoverable. For example, labeling DNA with fluorescent dyes, such as YOYO-1, was recently found to be elongating and twisting the native structure of DNA as well as changing its charging state (19, 20). Therefore, lots of efforts have been made in developing label-free techniques for single molecule detection, particularly, detection of DNA.

Atomic force microscopy (AFM) is a powerful label-free technique for detecting and manipulating single DNA molecules, however, it is limited by the scanning speed, and the scanning probe may also introduce perturbations to the target DNA (21–23). Another extensively investigated label-free detection approach is based on the conductance change induced by the target DNA in the electric circuit that is comprised of semiconductor nanomaterials when the DNA is hybridizing to its complementary probe which is covalently-linked to the surface of the detector or translocating through a nanopore fabricated in the device (24–26). These approaches were reported to achieve a high detection sensitivity. But the high cost and complexity in device fabrication puts a limit on its application. Although some recent reports introduced a convenient approach in fabricating glass nanopores in microfluidic devices for single DNA molecule detection (27, 28), it is not capable of distinguishing and detecting the sequence-specific targets; and like all the nanopore-based detection schemes, frequent clogging of the nanopore poses a big problem in practical applications.

In this report, we propose an IVT2H (in-vitro two-hybrid)-based detection strategy on the drop-based microfluidic platform to realize the label-free DNA detection on the single molecule level with a kHz interrogating rate. Specifically, the target DNA serves as the template for an in-vitro protein synthesis (IVPS), from which the synthesized protein (target protein) acts as the transcription factor to activate the expression of the reporter, a fluorescent protein in the droplet. Therefore, by detecting the signals from the reporter, we are able to detect the presence of the target DNA. As the picoliter volume of the drops effectively reduces the volume-proportional noises from impurities and solvent in the DNA solution (29, 30), and in-vitro transcription and translation realizes signal amplification, the detection sensitivity in our strategy successfully reaches the single molecule level, as demonstrated in the detection of a range of model DNA molecules.

## Materials and Methods

### Microfluidic device fabrication

Microfluidic devices were fabricated by patterning channels in polydimethylsiloxane (PDMS) using conventional soft lithography methods (31). Briefly, SU8-3010 photoresist (MicroChem Corp.) was spin-coated onto a 3” silicon wafer and patterned by UV exposure through a photolithography mask. After baking and development with SU-8 developer (propylene glycol methyl ether acetate; MicroChem Corp.), a 10-μm tall positive master of the device was formed on the silicon wafer. Then a 10:1 (w/w) mixture of Sylgard 184 silicone elastomer and curing agent (Dow Corning Corp), degassed under vacuum, was poured onto the master and cured at 65 °C for 2 hours. Afterwards, the structured PDMS replica was peeled from the master and inlet/outlet ports were punched out of the PDMS with a 0.75 mm-diameter biopsy punch (Harris Unicore). The PDMS replica was then washed with isopropanol, dried with pressurized air, and bonded to 50 × 75 mm glass slides using oxygen plasma treatment to form the device.

To enable the formation of aqueous-in-oil emulsions, the microfluidic channels must be made hydrophobic, which is routinely accomplished through Aquapel (PPG Industries) treatment. However, due to the existence of considerable biomolecule components in our aqueous phase, non-specific adsorption of these biomolecules to the PDMS channel will cause the severe wetting in the Aquapel treated device during the drop-making process. So instead of Aquapel, we passivated our drop-maker with a fluorosilane solution (1% (v/v) of 1H,1H,2H,2H-perfluorooctyl-trichlorosilane (Sigma-Aldrich) in Novec HFE-7500 oil (3M)) by flushing it into the device and immediately drying the channels with pressurized air. To stabilize our droplets against coalescence, we used EA surfactant donated by RainDance Technologies. The surfactant was dissolved in the fluorinated carrier oil Novec HFE-7500 at a concentration of 1.8% (w/w). PhD 2000 syringe pumps (Harvard Apparatus, Inc.) were used to infuse the fluids into the device.

### Preparation of the in-vitro synthesis reactions

All the DNA constructs used in this work were designed and synthesized as reported (32). The PURExpress^®^ in-vitro synthesis kit (New England Biolabs) was used for in-vitro protein synthesis. The manufacturer-recommended DNA concentration in single-expression bulk IVPS is 25-1000 ng/25 μl. To ensure the maximal production of the reporter protein in the multi-protein expressing IVPS, we controlled the amount of the target DNA slightly less than the reporter DNA. Specifically, reporter DNA was kept at 6 ng/μl while target DNA was diluted to be 3 ng/μl, 0.3 ng/μl, 0.24 ng/μl, 0.15 ng/μl and 0.03 ng/μl for the experiments at λ (average DNA molecule number per droplet) = 10, 1, 0.8, 0.5 and 0.1. We prepared the aqueous phase with 20 μl solution A (including necessary enzymes), 24 μl solution B’ (including small molecules and 300 ng reporter DNA), 2 μl target DNA and 4 μl Nuclease-free H2O, mixing them well for encapsulation into 10 μm-diameter droplets. For the detection of DNA *cro-ncoa1*, where partner DNA *er-ad* was introduced in the IVPS, 2 μl *er-ad* (3 ng/μl) was added and balanced by water to reach a total volume of 50 μl. All the reagents were handled on ice before emulsification to prevent the IVPS initiation in bulk.

### Fluorescence signal detection

As the drops passed through the laser spot in the 10 μm-wide channel (Fig. 1c), the light emitted from the fluorescent drops was captured by the objective and filtered through dichroic mirrors and the bandpass filter to the green photomultiplier tube (PMT, Hammamatsu) with the peak measurement band at 536/40. The output signal from PMT was acquired and processed by a computer-running LabView, executed on the field-programmable gate array (FPGA) card (National Instruments) at an acquisition rate of 100 kHz. A peak analysis algorithm was applied to analyze the data and identify the legal drops by interrogating the PMT output and time duration according to the set thresholds. The broad bursts that might have resulted from the coalesced drops and the spikes from the environmental interferences were discarded from the analysis. PMT gain was set at 0.36 for all the detection experiments unless otherwise specified.

**Fig. 1.**
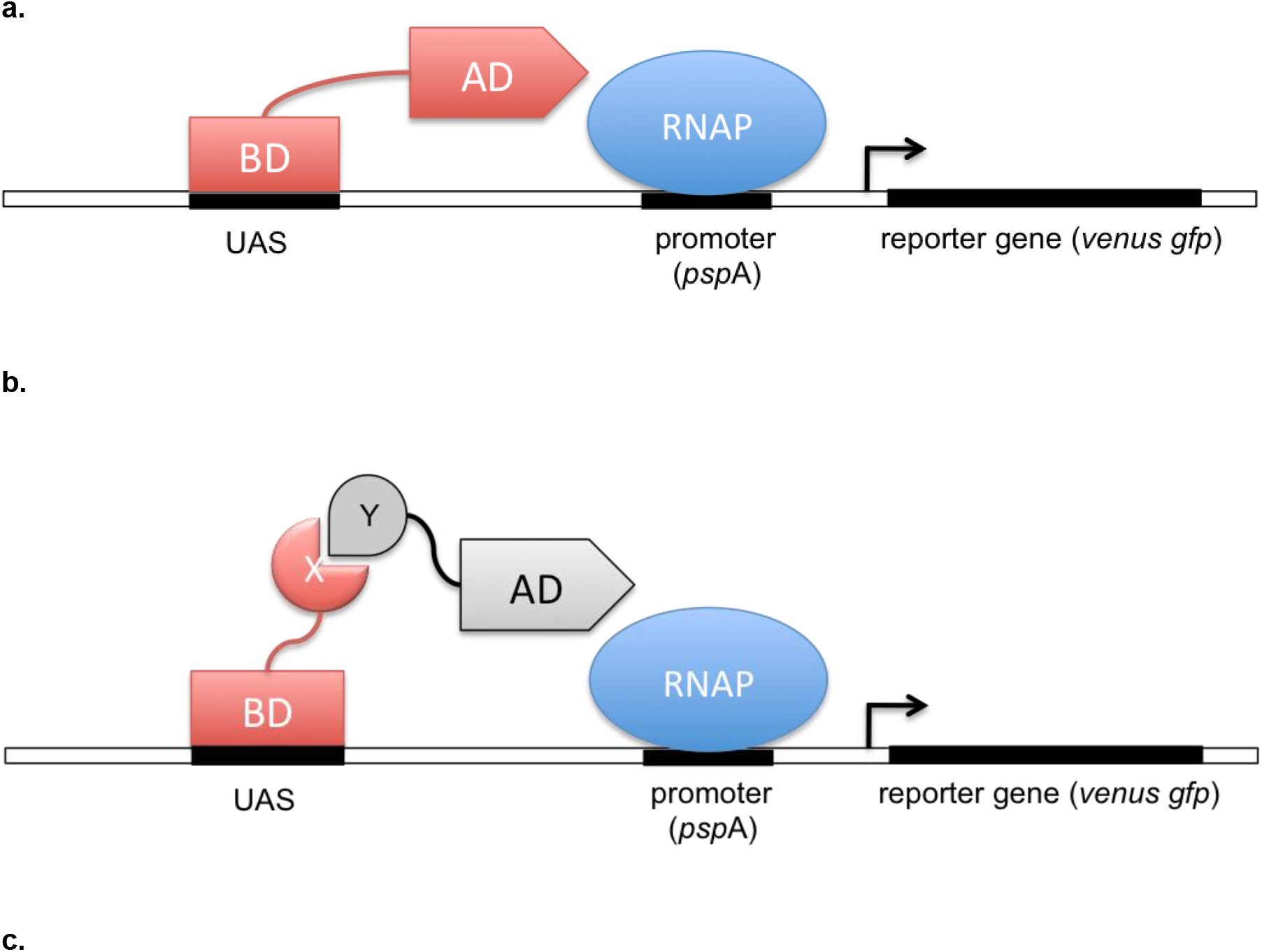
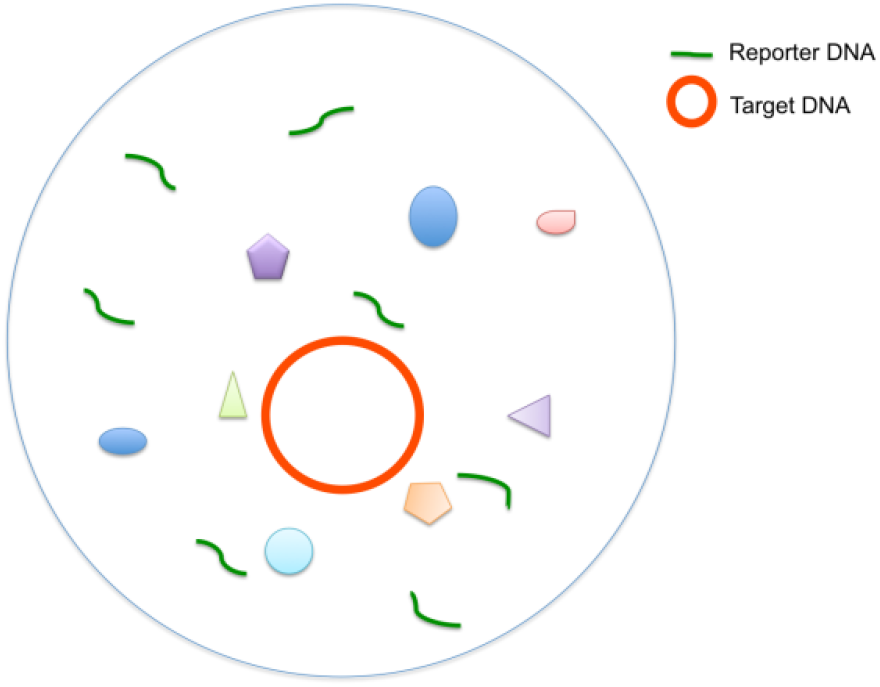
In-vitro two-hybrid-based detection strategy. (a) One-hybrid transcription regulation mechanism for DNA (encoding the protein in red) detection using DNA-protein interaction. Once the DNA binding domain (BD) binds to the upstream activating sequence (UAS) adjacent to the promoter (*psp*A) on the reporter gene, it activates the transcription initiation of the reporter gene by interacting with the promoter-bound E. coli RNA polymerase (RNAP) via the activation domain (AD), and hence triggers the translation of the fluorescent reporter gene (*venus gfp*). (b) Two-hybrid transcription regulation mechanism for DNA (coding for the protein in red) detection using protein (red)–protein (grey) interaction. An interacting protein pair X and Y are fused to BD and AD respectively, forming two hybrid proteins, which work together to activate the transcription. (c) Schematic of a drop containing the target DNA (in plasmid), reporter DNA (linear) and multicomponent IVPS kit. The components in IVPS kit are denoted with the irregular-shaped symbols in the picture.

## Results and Discussion

### DNA detection strategy on the drop-based microfluidics

The premise of our drop-based DNA detection strategy is the fact that one DNA codes for an increased number of its protein through repeated transcription and translation processes. This provides an initial signal transduction and amplification. Benefited from the isolated confinement provided by drops, the expressed protein can continue to trigger, as an activator, the expression of a second gene in the same droplet and therefore lead to a secondary signal transduction and amplification from the target DNA. And this avalanche signal transduction and amplification process can keep proceeding until a detectable signal level is reached. It is thus crucial to design a tight gene expression regulation mechanism that could reliably connect the target DNA with the reporter protein, via the intermediate protein products and their molecular interactions, for which we choose an adapted transcription regulation machinery from Escherichia coli (E.coli) σ^54^ promoter activating system and recapitulate it in the IVPS environment (32). In *E.coli*, the transcriptional activator PspF binds to the upstream activating sequence (UAS) with its DNA binding domain (BD) and activates the transcription with the activating domain (AD) via interacting with the inactive *pspA* (the promoter)-bound σ^54^ -RNAP complex (Fig. 1a). It has been verified in previous studies that replacing BD of PspF and its UAS with other DNA binding proteins and the recognition site does not affect the activation qualitatively (32), since BD and AD function independently. So we fused our target DNA with the DNA encoding AD of PspF (*ad*) to form the hybrid protein as the synthetic activator in our detection design. For the proteins that don’t specifically bind to a DNA sequence but specifically bind to a protein, their DNAs can be detected through a further modified two-hybrid strategy (Fig. 1b), where the target DNA x is fused to the DNA (*bd*) coding for BD; and its partner DNA y, whose protein has a specific binding to the target protein is fused to *ad*. Likewise, an intermediate RNA with both binding sequences to proteins X and Y can be introduced into the DNA-RNA hybrid for the detection of the DNAs whose protein interacts with RNA, instead of DNA or proteins. Since most of DNAs of interest in biological research have specific binding to other biomolecules for their physiological functions, this molecular interaction-based DNA detection strategy therefore has vast and practical applications. Additionally, the reported correlation between the transcription activation level and the molecular interaction strength in both cell-based and in-vitro two-hybrid experiments (32, 33) suggests that our drop-based detection system can be potentially applied to studying the unknown molecular interactions of biomolecules in a high-throughput approach, which has not been reported for conventional single molecule detection techniques.

We used venus-GFP as the reporter protein for its structural simplicity compared to widely used fluorescence-inducing proteins such as β-galactosidase. The adapted E. coli σ^54^ promoter activating system has been demonstrated working in bulk (32). And the cell-free protein synthesis in drops have previously been reported with single protein type such as EGFP, laccase, and β-galactosidase (34–36). However, in our system, multiple relevant proteins are to be expressed cascade in the same IVPS drop from a small number of DNAs, and therefore more details have to be taken into consideration in the experimental design for a maximal production of the reporter protein. Among many variables, the ratio of the DNA amount between the target gene and the reporter gene is especially critical, in that those two genes are under different promoters (a weak E. coli σ^54^ promoter for the reporter gene, and the strong T7 promoter for the target gene). This difference in transcription efficiency in a limited resource environment as in IVPS, would lead to a competition between the two genes for synthesizing resources such as enzymes, ATPs and small molecules, therefore reducing the target protein production. To avoid that, we fine-tuned the ratio of the DNA amount between the target gene and the reporter gene according to their translation efficiency and performed the IVTS in drop within the optimized ratio range.

### Target encapsulation and droplet IVPS

Fig. 2a gives the workflow of the microfluidic drop-based detection experiment. The mixture of the ice-cold IVPS kit and target DNA was introduced into the aqueous channel of the 10 μm drop–maker at a flow rate of 60 μl/h, and the fluorinated carrier oil Novec HFE-7500 (3M) with 1.8% (w/w) EA-surfactant (RainDance Technologies) was infused into the oil channel at a flow rate of 250 μl/h. Monodispersed drops of ~0.5 pL containing the IVPS-DNA mixture were formed at the flow-focusing junction (Length: 10 μm, Width: 10 μm, Height: 10 μm) of the microfluidic drop-maker (Fig. 2b) and collected into a 1 mL BD Luer-Lok™ syringe (BD Medical) for off-chip incubation. The number of the DNA molecules encapsulated in one droplet follows the Poisson distribution

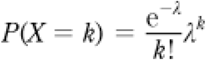

where P is the probability of having k DNA molecules in a drop when the average DNA molecule number per droplet is λ. The drops with target DNA inside would have contained the fluorescent protein venus GFP after a series of cell-free transcription and translation reactions during the 37 °C incubation. We incubated the emulsion in dark for ~6 hours to allow sufficient reactions in drops, after which the emulsion was reinjected into the 10 μm microfluidic detector for fluorescence signal detection (Fig. 2b). The emulsion was flowed into the detector at 20 μl/h and spaced out with the fluorinated carrier oil Novec HFE-7500 (200 μl/h) at the arrow-shaped junction before entering the detection region where the single-file lined drops were scanned by the 488 nm laser (Newport) for analysis.

**Fig. 2.**
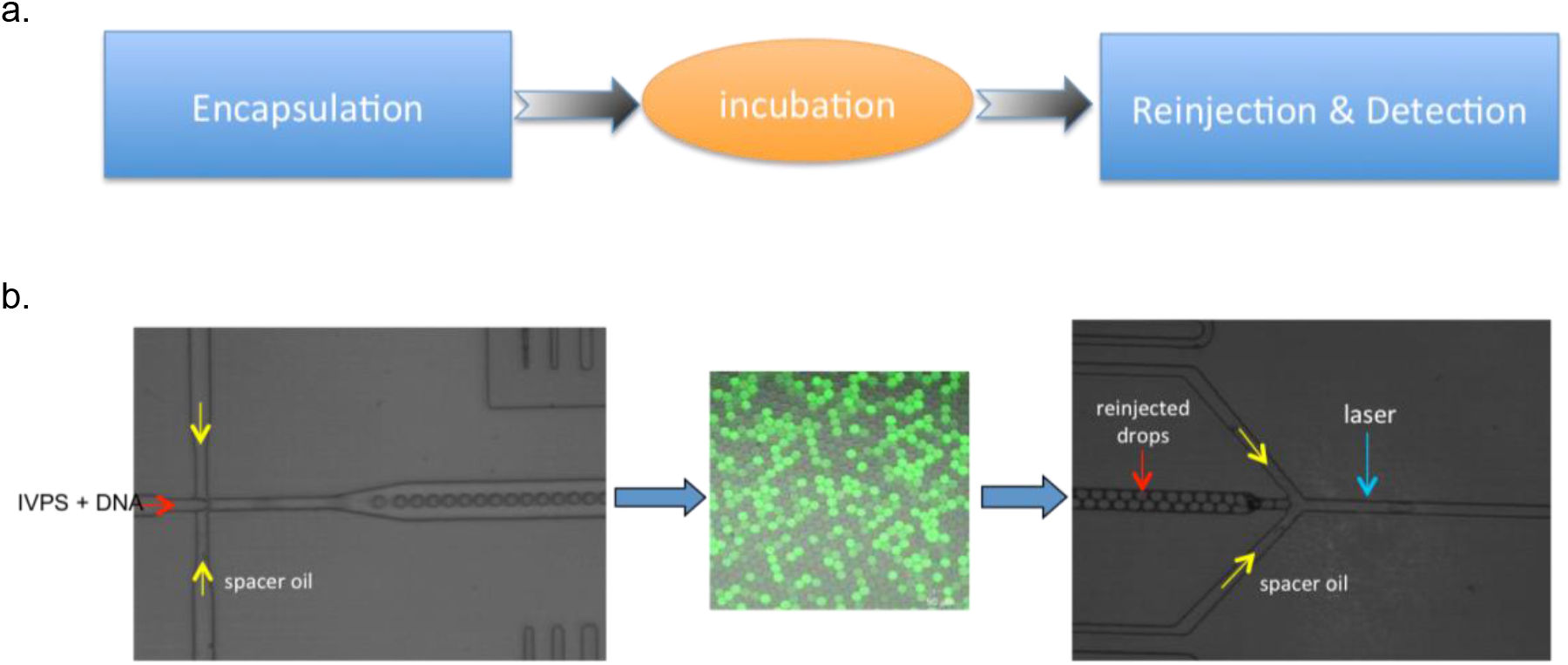
DNA detection design with drop-based microfluidics. (a) Schematic diagram of the experimental steps for the microfluidic drop-based DNA detection. Off-chip operation is denoted in orange. (b) Representative pictures of the post-incubated emulsion (middle, overlay of the fluorescence image and bright field image), drop-maker (left) and detector (right). Emulsion flows from left to right in the detector. Channel depth: 10 μm. Narrowest channel width: 10 μm.

### Detection of the single DNA molecules

#### 1. DNA detection via protein-DNA interaction

For its small size (~20 amino acids) and high binding affinity (Kd~pM) (37) to the consensus DNA binding site, we chose phage λ protein Cro as the model target protein to replace the BD of PspF for proof of principle. As illustrated in Fig. 1a, we aimed to detect the DNA *cro-ad* that encodes the protein Cro-AD in the droplet, based on the interaction of Cro and the consensus binding site on the reporter gene construct.

To characterize the drop-based detection system, we first measured the fluorescence from the “empty” drops, in which no target DNA was included. Time traces of fluorescence intensity were recorded (in Fig. 3a). Each peak above the baseline level corresponds to a trajectory of the fluorescent drops captured by the PMT detector when it was passing through the detection window. The peak width indicates the drop size, and the frequency of the peaks corresponds to the occurrence of the fluorescent drops at a given flow rate. In the experiment without target DNA in the drops, a moderate fluorescence level was observed in each droplet, as represented in a 10 ms snapshot of the fluorescence time traces (Fig. 3a). This background fluorescence indicates the weak expression of the reporter protein in the absence of the transcription activator. Its consistent intensity among drops served as the droplet indicator and thus saved the extra step of incorporating indicator dyes in the droplet as routinely used in drop experiments (30, 35, 38). When as many as 10 DNA molecules (λ = 10) were encapsulated into the drops, a similar uniform fluorescence level over the majority of the drops was observed in the time trace plot, with a 20 fold increase in intensity. A close examination on the drop number distribution over the fluorescence intensity at λ = 10 uncovered a small peak at the background fluorescence level (Fig. Sb), indicating the existence of a small population of the “empty” drops, which agrees well with the Poisson distribution (P_(X=0, λ=10)_ = 4.54E-5). A prominent difference rose up in the single-molecule detection experiments, for example, when the target DNA was diluted to a concentration of λ = 1. The fluorescence intensity no longer appeared uniform among drops (Fig. 3c). It is more clearly displayed in the fluorescence distribution histogram (Fig. 3d) where a significant portion of drops (~0.359) are “empty”, in excellent agreement with Poisson distribution (P_(X=0, λ=1)_ = 0.37), and thus demonstrates the successful detection of the single-occupancy drops. Experiments with more diluted DNA (λ < 1, data not shown) further verified the system’s capability in detecting single *cro-ad* DNA molecules in drops. The fluorescence intensity in the single-DNA drops can be obtained from the pivot point (most probable occupancy event) in the drop fluorescence distribution curve (λ ≤ 1).

**Fig. 3.**
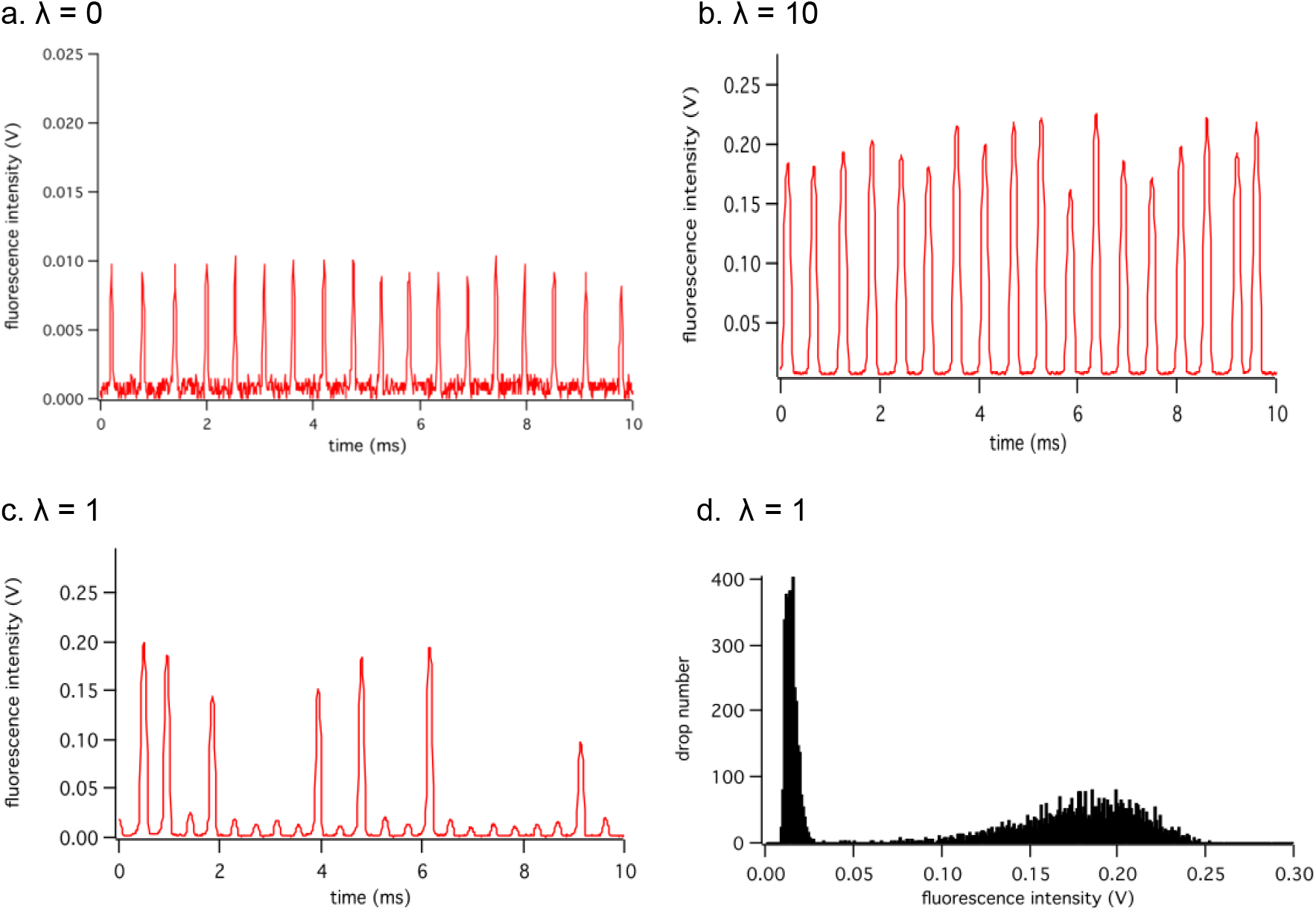
Detection of the DNA *cro-ad* in drops. Time traces of the fluorescence intensity for different *cro-ad* concentrations: (a) λ = 0, (b) λ = 10 and (c) λ = 1. Every peak above the baseline corresponds to a droplet. The height and width of the peak provide the information on the fluorescence and the size of the drop. The frequency of the peaks can be adjusted by the flow rates of the reinjected emulsion and spacer oil. (d) Drop fluorescence distribution obtained from the experiment at λ = 1. Drops were identified from the time trace data with the peak analysis algorithm executed on the FPGA.

As demonstrated above, for proteins like Cro-AD that have a high binding affinity to the consensus sequence, our system can detect its DNA at the single molecule level unambiguously. To test the system’s tolerance to weaker binding affinities, we performed a series of detection experiments on the targets that encode zinc finger proteins as the synthesized BD. Zinc finger is another small (23-28 amino acids) DNA-binding protein that is widely used in protein engineering as a preferred scaffold for creating customized DNA-binding domains with improved binding affinity and specificity (39–41). It binds to the DNA recognition site through multiple finger-like motifs coordinated by zinc ions. Via making changes to the original fingers or adding more fingers, higher affinity or specificity can be obtained in the zinc finger variants. For example, the reported binding affinity Kd of the wild type zinc finger Zif268 (Zif) to its wild type binding sequence is in the range of 10-500 pM (42, 43), much lower than that of Cro to its consensus sequence. Correspondingly, our experimental results with *zif-ad* as the target DNA showed no discernible difference in fluorescence intensity between the “empty” drops and the drops containing *zif-ad* at λ = 1 (Fig. 4a), suggesting that the transcription of the reporter gene was not effectively activated, due to the weak interaction between the BD Zif and the cognate sequence. Whereas a variant Zif//NRE screened by J.-S. Kim and C. O. Pabo (43) was claimed to have a much higher binding affinity (<1 pM) to its binding site. In our detection experiments with *zn-ad* (encoding Zif//NRE-AD), we were able to observe the two populations of “empty” and occupied drops at λ = 1 (Fig. 4b), indicating the existence of the fluorescence from the target-containing drops with successful activation on transcription, hence suggesting a stronger interaction of Zif//NRE to its binding site as reported. And the peak point in the occupied population corresponds to the most probable drops whose fluorescence, in the case of λ = 1, indicates the fluorescence from the single DNA-bearing drops. In addition to the successful detection of the target DNA at the single molecule level as discussed above, the stronger protein-DNA interaction from Zif//NRE-UASzn compared to Zif-UASz in our experiments is in good accordance with the reported binding affinities for those two protein-DNA interaction pairs, which verifies the positive correlation between the gene expression level and the interaction strength of its transcription activator and UAS in our drop IVPS-based two-hybrid system. It supports the similar finding in bacteria-two-hybrid system about the correlation between the transcription activity and the interaction strength of the activator protein pair (40), and enables the system to probe unknown molecular interactions in addition to the DNA detection.

**Fig. 4.**
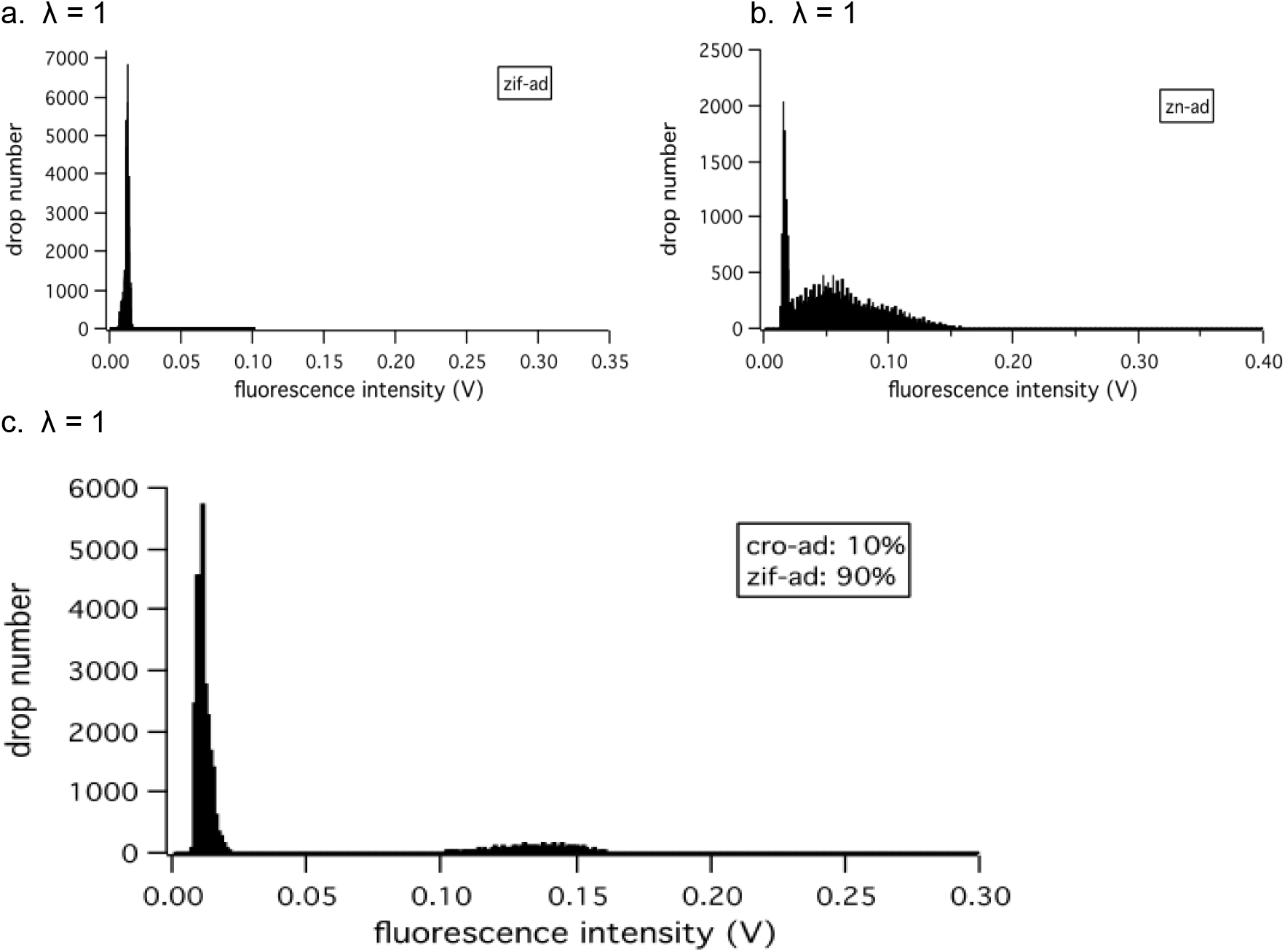
Detection of the single molecule DNA via weak protein-DNA interactions. Drop fluorescence intensity histograms obtained from the experiments with different target DNA templates at λ = 1: (a) *zif-ad*, which encodes a protein with the wild type Zif268 as the DNA binding domain. (b) *zn-ad*, which encodes a protein with the variant Zif//NRE as the DNA binding domain. PMT gain was raised to 0.5 for clarity. (c) mixture of *zif-ad* and *cro-ad* (λ_total_ = 1) with the ratio of 9:1.

Aforementioned detection experiments were all performed on the single-type target in an IVPS droplet. In fact, our system could also detect the target in a multiple-DNA cocktail, as exemplified in Fig. 4c, where the target DNA *cro-ad* and control *zif-ad* (as to the consensus sequence of Cro) were mixed at 1:10 with a total λ of 1. The population containing the target was identified as the small hump in the drop fluorescence histogram. It thus encouraged us to try detecting more types of DNA based on more complicated molecular interactions.

#### 2. DNA detection via protein-protein interaction

So far, we have successfully detected the single target DNA based on different protein-DNA interactions in the transcription regulating mechanism. However, not all the proteins have a fair binding affinity with UAS to be the activator in the protein-DNA interaction-based transcription regulation system. To expand the application of the assay, we challenged our system in detecting single DNA molecules based on protein-protein interactions (Fig. 1b), where the activator BD-X synthesized from the target DNA works in concert with a partner protein Y-AD to initiate the transcription of the reporter gene. If there is enough strength of the specific interaction between X domain and Y domain, the protein pair will function as one activator to initiate the expression of the reporter protein, thus producing the detectable fluorescence signals in the droplet. As a demonstration, we used the interaction between human estrogen receptor α (ER) and nuclear receptor coactivator 1 (NCOA1) under the modulation of the small molecule 17β-estradiol (E2) to detect the target DNA *bd-ncoa1* at the single molecule level.

The partner protein ER-AD was synthesized in IVPS reaction along with the target protein BD-NCOA1 to trigger the synthesis of the reporter protein venus-GFP in the droplet. As discussed above, the amount of the DNA molecules for the partner protein and reporter protein as well as the modulator molecules were optimized for the maximal production of the reporter protein in IVPS. Target DNA was coencapsulated with the partner DNA (λ = 10), E2 (1 μM) and IVPS kit into the 10 μm-diameter droplet before the reaction-inducing incubation off-chip. We measured the fluorescence of the reinjected droplets at ~2 kHz for a series of target DNA concentrations (Fig. 5).

**Fig. 5.**
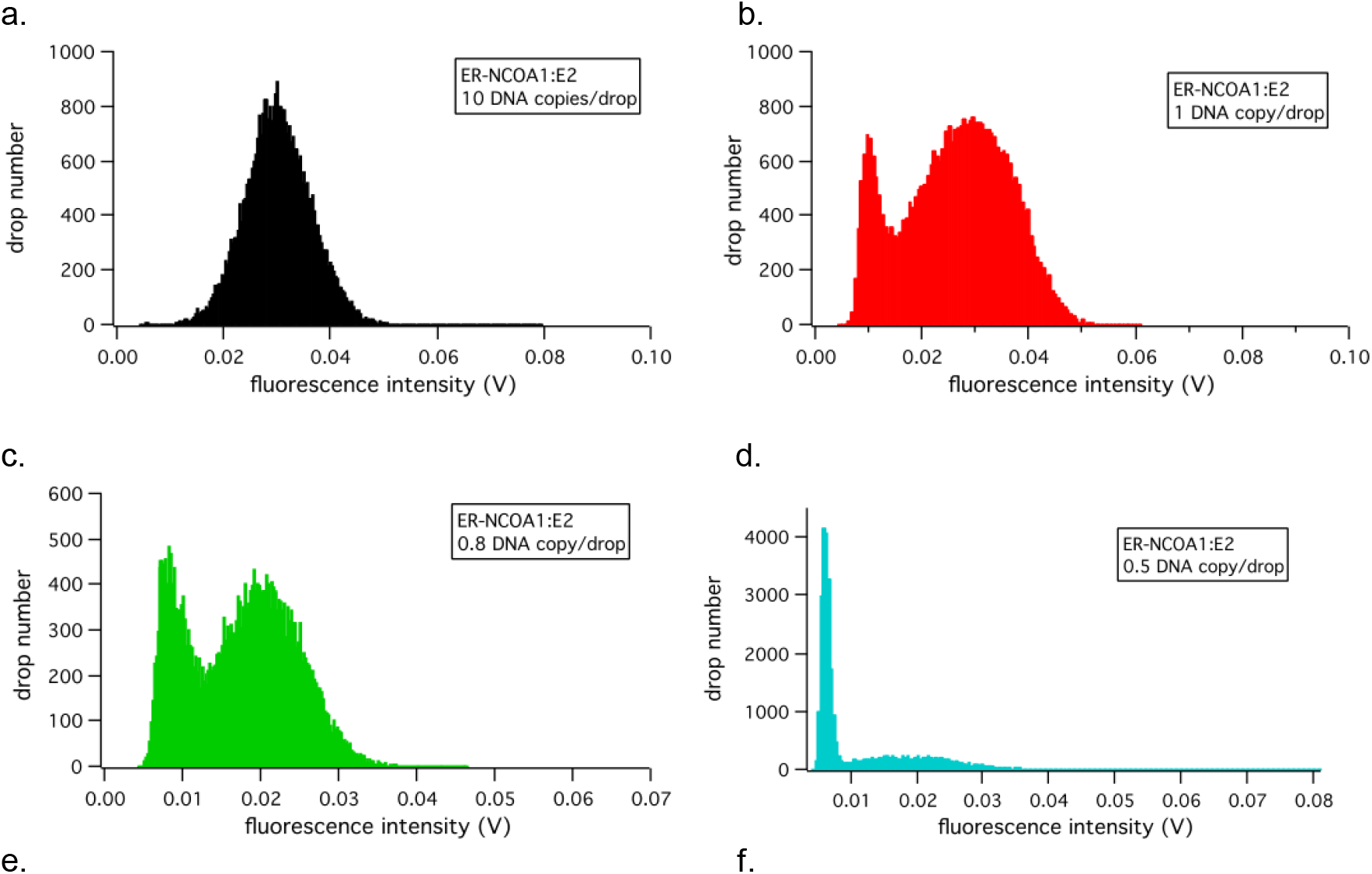
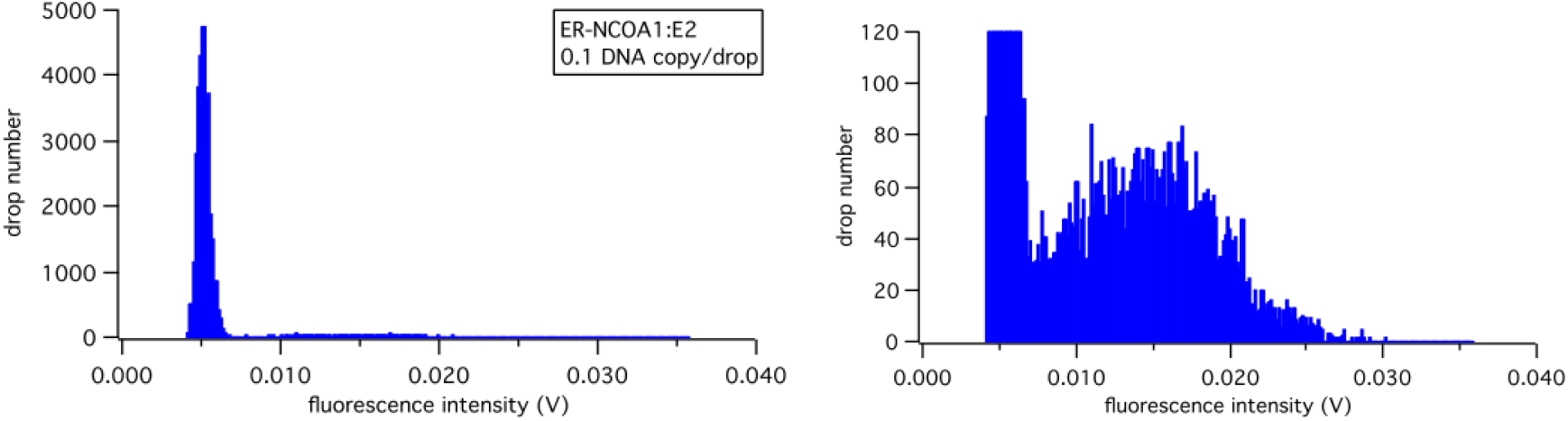
Detection of the DNA *bd-ncoa1* in drops. Drop fluorescence intensity histograms for different *bd-ncoa1* concentrations: (a) λ = 10, (b) λ = 1, (c) λ = 0.8, (d) λ = 0.5, (e) λ = 0.1. (f) is a zoom-in of (e) on the low y-value region for a clear view on the high-intensity population. The gene encoding the partner protein ER-AD was kept at λ = 10, and E2 was kept at 1 uM in all the *bd-ncoa1* detecting experiments.

At λ = 10 (Fig. 5a), the majority of the drops showed a fluorescence signal above the background level, indicating a predominant DNA-occupancy among the drops. The presence of the small population at the background fluorescence level agreed well with the Poisson distribution. In the experiments of λ ≤ 1 (Fig. 5b-5f), two populations of the drops became increasingly discernible in the drop fluorescence histogram, with the lower fluorescence drops gradually outnumbering the higher fluorescence ones as λ decreased, indicating the increase in “empty” drop number and single-occupancy occurrence. The consistence of the experimental drop distribution with Poisson distribution verified the detection of the single *bd-ncoa1* molecule. It not only demonstrates the robustness and sensitivity of our detection system in its ability to incorporate a complex multicomponent in-vitro protein synthesis, but also expands the range of the detectable DNA by introducing a partner protein to allow for a variety of molecular interactions to be utilized for the detection. Here we used BD-NCOA1 as the target protein; similarly ER-AD can also be set as the target protein if the optimized amount of *bd-ncoa1* is fixed. Based on the same principle, proteins that have interactions with other biomolecules, for example, RNA, can be the target protein as well for the detection of its DNA, where RNA can be tailored as a bridge between the activator pair to initiate the expression of the reporter protein.

In the *bd-ncoa1* detection experiments, we noticed a subtle λ-dependent (λ ≤ 1) left-shift trend of the fluorescence intensity in both the single-occupancy and the “empty” drops in the drop fluorescence histogram. In other words, at the single molecule level, as the target DNA concentration decreased, the fluorescence level from “empty” drops and the drops containing the single target DNA molecule both decreased in a subtle way. However, no similar changes were observed in the λ > 1 experiments. The constant fluorescence level in the λ > 1 experiments can be explained as the IVPS reached its plateau for the incorporated synthesis reactions and therefore any increase in the target DNA amount would not cause the increase in the production of the reporter protein. But interpretation to the intensity shift at the single molecule level is not that straightforward. Since the fluorescence was from the fluorescent protein that has a large molecular weight compared to the small fluorophore molecules, the chances of the fluorophore exchange between drops due to the concentration gradient observed in experiments with fluorescent dyes (36, 44, 45) are slim. Nevertheless, this subtle shift of both signal and background levels in the diluted DNA experiments does not affect our ability in detecting the single DNA molecule.

## Conclusion

To our knowledge, it is the first time the label-free single molecule DNA detection has been demonstrated inside a microfluidic droplet. The system detects target DNA through monitoring the interaction of its protein with other biomolecules. Since the detection is based on its intracellular bioactivity, there are no concerns of potential disturbance from external agents to the target. The detection strategy utilized here also renders this system the capability in exploring the molecular interactions from poorly understood or mutated biomolecules in diseased cells, for example, the detection of the mutated protein in cancer cells by monitoring its failed recognition of the binding sequence. Given its ultrahigh throughput interrogation rate, this drop-based microfluidic detection system might potentially benefit the areas such as rapid disease diagnosis, drug screening and protein engineering.

## Acknowledgements

We thank Dr. S. Chong at New England Biolabs, *Inc*. for generously providing IVT reagents and DNA constructs. We are grateful for technical assistance from Dr. Linas Mazutis with microfluidic devices.

This work was supported by the National Science Foundation (DMR-1310266) and the Harvard Materials Research Science and Engineering Center (DMR-1420570).

