## Supplemental Figure for "PCR-free, label-free detection of sequence-specific DNA with single-molecule sensitivity using *in vitro* N-hybrid system in microfluidic drops"

**Supplementary Information**

a. λ = 0

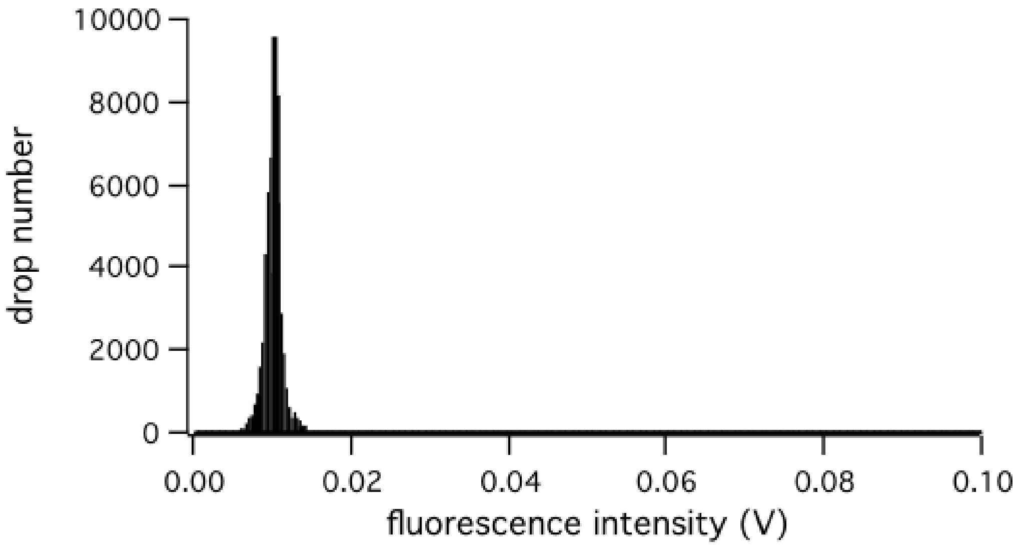


b. λ = 10

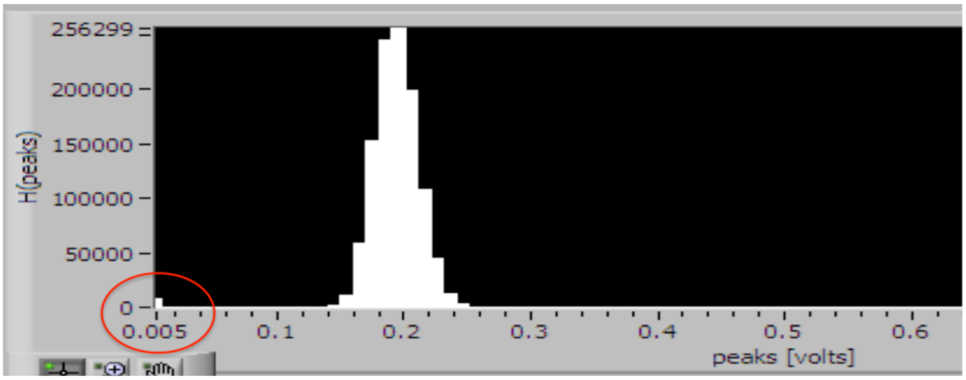


**Fig. S Drop fluorescence intensity histogram from *cro-ad* detecting experiments:** (a) λ = 0, (b) λ = 10. The red circle in (b) highlights the population with background fluorescence.
